# Inference of candidate germline mutator loci in humans from genome-wide haplotype data

**DOI:** 10.1101/089623

**Authors:** Cathal Seoighe, Aylwyn Scally

## Abstract

The rate of germline mutation varies widely between species but little is known about the extent of variation in the germline mutation rate between individuals of the same species. Here we demonstrate that an allele that increases the rate of germline mutation can result in a distinctive signature in the genomic region linked to the affected locus, characterized by a number of haplotypes with a locally high proportion of derived alleles, against a background of haplotypes carrying a typical proportion of derived alleles. We searched for this signature in human haplotype data from phase 3 of the 1000 Genomes Project and report a number of candidate mutator loci, several of which are located close to or within genes involved in DNA repair or the DNA damage response. To investigate whether mutator alleles remained active at any of these loci, we used *de novo* mutation counts from human parent-offspring trios in the 1000 Genomes and Genome of the Netherlands cohorts, looking for an elevated number of *de novo* mutations in the offspring of parents carrying a candidate mutator haplotype at each of these loci. We found some support for two of the candidate loci, including one locus just upstream of the *BRSK2* gene, which is expressed in the testis and has been reported to be involved in the response to DNA damage.

**Author Summary:** Each time a genome is replicated there is the possibility of error resulting in the incorporation of an incorrect base or bases in the genome sequence. When these errors occur in cells that lead to the production of gametes they can be incorporated into the germline. Such germline mutations are the basis of evolutionary change; however, to date there has been little attempt to quantify the extent of genetic variation in human populations in the rate at which they occur. This is particularly important because new spontaneous mutations are thought to make an important contribution to many human diseases. Here we present a new way to identify genetic loci that may be associated with an elevated rate of germline mutation and report the application of this method to data from a large number of human genomes, generated by the 1000 Genomes Project. Several of the candidate loci we report are in or near genes involved in DNA repair and some were supported by direct measurement of the mutation rate obtained from parent-offspring trios.

## Introduction

The rate of germline mutation is a key parameter in molecular evolution and population genetics. As the ultimate source of genetic novelty, germline mutations provide the raw material on which selection acts and the basis for genetic drift over time. Mutation rates are known to differ substantially between species [1], and in eukaryotes, the single nucleotide mutation rate, fundamental to many demographic and evolutionary analyses, ranges over two orders of magnitude [2]. Methods to estimate the rate of *de novo* mutation predate the knowledge that DNA carries hereditary information. If the frequency of a deleterious allele is in mutation-selection balance, the rate at which deleterious alleles are removed through selection is equal to the rate at which novel alleles arise through mutation. This idea was used by Haldane to provide an indirect estimate of the rate of spontaneous haemophilia from prevalence estimates [3]. Subsequent estimates incorporated knowledge of the physical size of the locus (or more precisely, the number of target nucleotides giving rise to the phenotype of interest) to obtain per-base and per-generation mutation rate estimates (e.g. [4, 5]).

More recently, whole genome resequencing methods have been used to obtain direct measurements of the human mutation rate from parent-offspring trios [6–9]. As well as enabling measurement of the genome-wide mutation rate, these studies have opened up the possibility of investigating factors contributing to variation in mutation rate between individuals. A study of 78 Icelandic parent-offspring trios estimated that 97% of variation in the number of *de novo* mutations called in offspring could be explained by the age of the father [8], and other whole-genome studies have shown similar, albeit weaker, relationships between parental age and germline mutation rate [10–12].

Genetic and environmental factors may also influence the rate of germline mutation [10, 13], and a number of recent studies point to differences in the germline mutation spectrum between human populations [13, 17, 18], consistent with a genetic contribution to variation in germline mutation. Indeed, several cancer-associated germline human mutations are known to affect genes involved in DNA proofreading and mismatch repair, increasing cancer risk through an elevated rate of somatic mutation [14, 15]. For example, Lynch syndrome, which is associated with a very high lifetime risk of development of cancer of the colon and several other organs, results from germline mutations in a number of mismatch repair genes [16]. Although the effects of such mutations on the rate of germline mutation in human have not yet been determined, deficiency in mismatch repair is known to result in an elevated per generation mutation rate in yeast [10].

Here we investigate the feasibility of detecting genetic polymorphism associated with increased germline mutation rate by looking for an increase in the number of derived alleles in haplotypes carrying the mutator allele. Using simulation, we show that a mutator allele can result in a localized peak in the numbers of derived alleles in a subset of haplotypes, against a background of haplotypes with typical numbers of derived alleles. This is because in the region linked to the mutator allele, haplotypes containing it are always subject to the elevated mutation rate, whereas other haplotypes are only affected when they occur together with the mutator allele in a heterozygote, and this will occur only rarely if the mutator allele is rare. Detecting this pattern depends on persistence of the mutator allele for a large number of generations, and we discuss the likelihood of this given estimates of the selective disadvantage of a mutator allele. We search for this pattern in data from the 1000 Genomes Project (G1K) and report and characterise a number of candidate mutator loci. For a subset of the candidate loci the highly derived haplotypes, characteristic of a mutator allele, were found among parents of two human trio datasets obtained from the Genome of the Netherlands (GoNL) [19] and G1K [20] projects and in each case we tested for an elevated number of *de novo* mutations in the offspring of parents carrying a putative mutator haplotype.

## Results and Discussion

We hypothesized that mutations that increase the rate of germline mutation in human populations may be detectable from genome-wide polymorphism data. Consider a population with a genetic variant at a locus that increases the germline mutation rate by a factor *ϕ* (in heterozygotes). If the variant arose *g* generations before the present, we can estimate the number of additional derived alleles, per site, that are expected in haplotypes containing the mutator allele over haplotypes with the wild-type allele as *gμ*(*ϕ* 1), where *μ* is the wild-type mutation rate. Let *ρ* be the rate of recombination per generation, which, for simplicity, we assume is uniform across the genome. The lengths of the genomic segments to the right and left of the mutator locus that have been free from recombination for *g* generations are then exponentially distributed with mean 
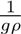
 and therefore the segment around the mutator locus that has been free from recombination since the mutator variant arose has an expected length of 
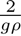
. The expected number of additional mutations in a segment of this length is 
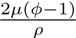
. Note that this is independent of *g*, the age of the mutator allele. Taking *μ* = 1.2×10^−8^ [8], *ρ* = 1.26×10^−8^ [21], gives an additional 1.9(*ϕ*−1) mutations expected in a segment of mean length.

Whether it is feasible to detect a local peak in the density of derived alleles clearly depends on *ϕ*, the affect of the mutator allele on the germline mutation rate. It also depends on the background number of derived alleles in haplotypes unaffected by the mutator allele. This, in turn, depends on *g* through the mean segment length. For example, if *ϕ* is 5 and *g* is 20,000 generations, then the expected number of additional mutations in a segment of expected length is 7.6 and the mean segment length is approximately 8 Kb (this is likely an underestimate since it assumes uniform recombination). Assuming a pairwise heterozygosity rate in humans of 0.001 per bp [22] implies that we expect approximately 8 derived alleles per background haplotype in an 8 Kb segment. Although we expect a relatively small number of additional mutations in an affected haplotype, the signal of an excess in the number of derived alleles may be carried by several related haplotypes, such that the total number of unique excess mutations among the affected haplotypes may be much greater, potentially resulting in greater power to detect the affected loci.

### Persistence of a mutator allele

Because the number of additional derived alleles is independent of the time since the mutator allele arose, but the segment length is not, the power to detect a mutator allele above the background is greater for alleles that arose a large number of generations before the present. Given that the mutator allele is also required to be associated with a large effect on the germline mutation rate, it is necessary to consider whether such an allele could persist for a large number of generations despite the reduction in fitness that will result from an increased germline mutation rate. Although the increased burden of *de novo* mutations associated with the mutator allele is likely to reduce the fitness of individuals carrying it, this reduction may be relatively small [23], and the per generation germline mutation rate is capable of sustaining substantial variation in animals, for example ranging widely across mammalian species [2]. Bear in mind also that *de novo* mutations will remain linked with the mutator allele only for an average of two generations before recombining away [2]. Indeed, a mutator allele, even at relatively low frequency, may contribute a large number of novel mutations, exerting a disproportionate influence on the evolution of the population as a whole while it persists.

The selective disadvantage associated with a (heterozygous) allele that increases the germline mutation rate can be approximated as 2*s_d_*Δ*U* [2], where *s_d_* is the mean selective disadvantage of a heterozygous deleterious mutation, *U* is the genomic deleterious mutation rate and Δ*U* is the change in *U* resulting from the mutator allele. Thus given a value for *U*, a mutator allele that increases the germline mutation rate by a factor *ϕ* will be associated with a selective coefficient (for heterozygotes) of *s* = 2*s_d_*(*ϕ* − 1)*U*.

The parameters *s_d_* and *U* are not straightforward to estimate. Lynch [2] provides values for multicellular eukaryotes from the literature that range over orders of magnitude from 10^−3^ – 10^−2^ for *s_d_* and from 0.01 – 1 for *U*. For values at the lower ends of these ranges, even a mutator allele associated with a large value of *ϕ* would have a relatively small effect on fitness. For example, *ϕ* = 5 gives *s* = −0.00008, which is in the region in which genetic drift begins to dominate over selection for an effective population size on the order of 10^4^. In this case a strong mutator allele could persist in a population for a large number of generations, sufficient to have an impact on the number of mutations that accumulate in the region linked to the mutator allele. By contrast, values of *s_d_* and *U* at the higher ends of these ranges would give *s* = −0.08, indicating strong selection against a mutator allele. Such an allele would not persist for long and would rapidly reduce in frequency in an organism with even a moderate effective population size.

Estimates of *U* in particular genomic regions have been made by comparing between-species sequence divergence within these regions with divergence in putatively neutral regions (such as fourfold degenerate coding sites, pseudogenes or inactive transposable elements). Using this approach, Keightley [24] estimated a relatively high value of 2.2 deleterious mutations per generation across the whole genome in humans. There are caveats associated with this estimate (such as the assumption that genomic mutation rates are the same across sequence categories and possible bias from alignment in calculating genome-wide divergence), but it does seem likely that *U* is not substantially less than one.

The mean selective disadvantage is more difficult to estimate, due in part to the confounding influence of demographic factors. Boyko *et al.* [25] estimated values of around −0.03 for the mean selective disadvantage of heterozygous amino acid changing mutations in African Americans. However, this is likely to be unrepresentative of mutations genome wide and heavily influenced by strongly deleterious mutations. Also, although Boyko *et al.* [25] assumed a codominant selection model, many of these mutations are likely to be recessive and thus may not be deleterious when they arise *de novo* (as they are almost certain to be heterozygous). For *Drosophila* García-Dorado and Caballero [26] estimated a mean of 0.1 for the coefficient of dominance, *h*, defined such that the fitnesses of heterozygotes and homozygous mutants are 1 + *hs* and 1 + *s* respectively, and *h* = 0.5 corresponds to codominance. The best-fitting model of Boyko *et al.* [25] for the distribution of selective effects of amino acid changing mutations included a normally distributed component of weakly deleterious mutations with mean of −0.0002, as well as a point mass representing strongly deleterious mutations. Consistent with this, Do *et al.* [27] have shown that the absence of differential removal of deleterious mutations from the genomes of Africans and non-Africans implies that selection coefficients acting on non synonymous substitutions are either strong (*s* < 0.004) or very weak (*s* > − 0.0004). In our context it may be possible to neglect strongly deleterious mutations on the grounds that they are typically recessive and rare outside protein-coding regions, where most *de novo* mutations occur. Mean selection coefficients for mutations outside protein coding regions are likely to be much smaller. If we set *s_d_* = − 0.0001, *U* = 1, *ϕ* = 5 and set *h* = 0.1 (to allow for the fact that even weakly deleterious mutations may have a coefficient of dominance well below 0.5), the selective disadvantage of the mutator allele in heterozygotes is approximately −0.0001. Such a mutation is affected by selection, but in a species with an effective population size similar to that of humans it can persist for a large number of generations by chance, particularly if it attains a high initial frequency due to a population bottleneck. The expected number of generations for which a mutation persists, under codominant selection, can be calculated [28]. For example a mutation with initial frequency of 0.2 and a selective coefficient of −0.0002 (−0.0001 in heterozygotes) has an expected time to extinction of 13,000 generations.

### Proof of concept simulations

We used simulation to investigate the effect of a mutator allele on the numbers of derived alleles in the region in linkage disequilibrium with it. We considered a genomic region of 100 Kb, with a mutator locus *x* at 50 Kb, and implemented simulations in a coalescent framework, with recombination and selection (see Materials and Methods for details). As expected, where mutations with a large impact on the germline mutation rate remained polymorphic in the population for a sufficient time, the genomic region close to the mutator locus was frequently found to exhibit a peak in the number of derived alleles, with this peak shared by a subset of haplotypes and the remaining haplotypes having a typical number of derived alleles across the region. An example of a simulation that exhibits such a peak is shown in Fig 1, and the results of ten simulation runs are provided as supplementary information Fig. S1. Whether a simulation displays this characteristic signature of a mutator locus depends on the history of recombination and coalescence in the surrounding genomic region. For example, recombination events close to the locus can disrupt linkage, eroding the increase in the number of derived alleles associated with the mutator locus. For the simulation shown in Fig 1 the mutator allele arose 20,000 generations before the present and resulted in a five-fold increase in the germline mutation rate in heterozygotes (and a ten-fold increase in homozygotes).

**Fig 1.**
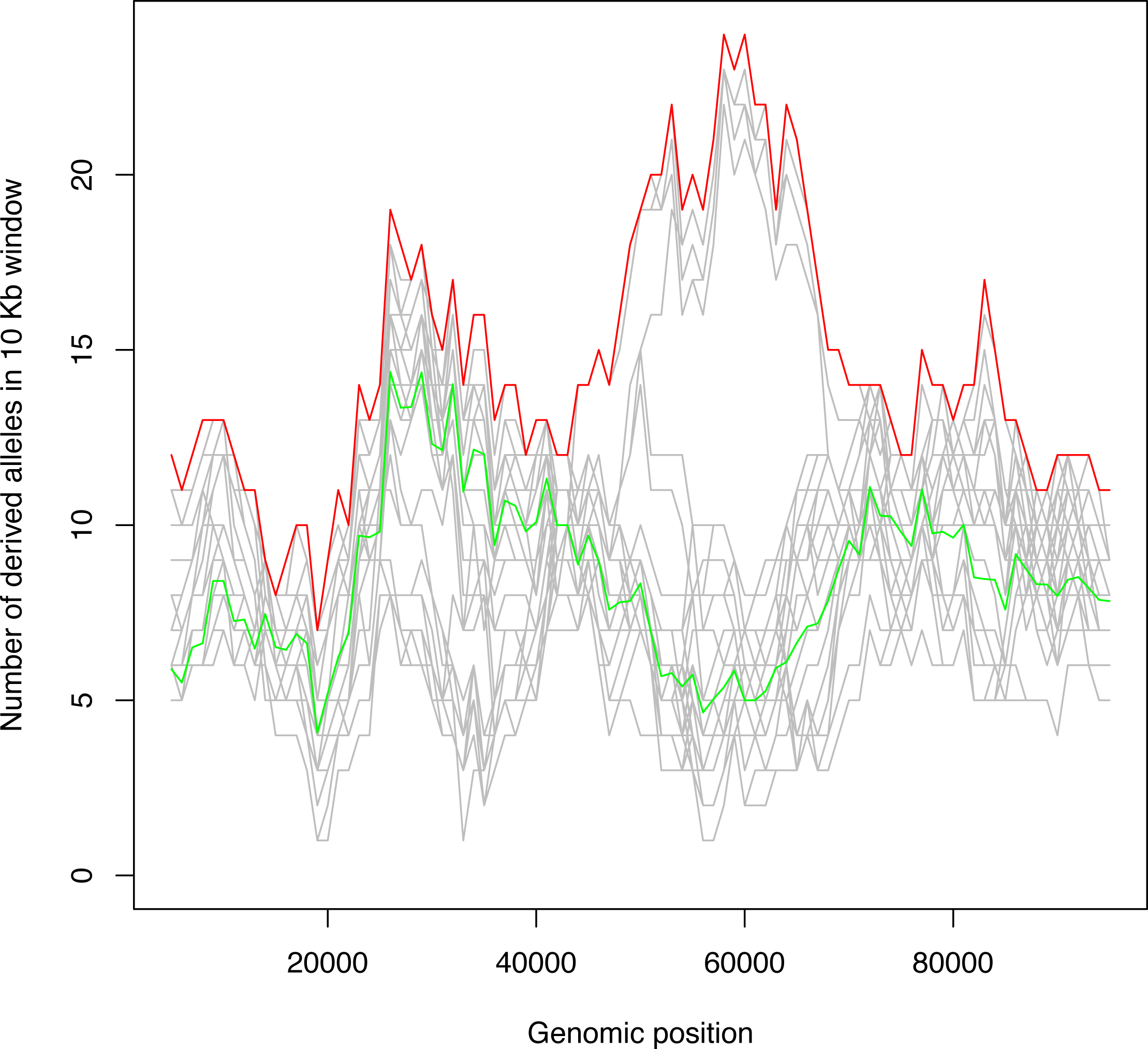
Simulation study. An example of simulated data showing a subset of haplotypes with a peak in the number of derived alleles in a 10 Kb sliding window. A region of 100 Kb was simulated over 40,000 generations using a coalescent approach with recombination. In this example a mutator allele with *ϕ* = 5 was introduced 20,000 generations before the present and was assumed to be weakly deleterious, with a selective coefficient of −0.0002. The red and green lines show the maximum and trimmed mean number of derived alleles in the window. Individual sampled haplotypes are shown in grey.

### Genome scan for candidate mutator loci in humans

To search for human loci that may have carried mutator alleles, we obtained data from phase 3 of the G1K project [20]. The data consisted of inferred haplotypes for a total of 2504 individuals in 26 populations, along with putative ancestral alleles for each variable site inferred from Ensembl Compara release 59 [29]. For each haplotype in a population we counted the number of derived alleles in a sliding window of size 10 Kb along the genome. In each window we then calculated the maximum number of derived alleles in the window and the interquartile mean number of derived alleles across all haplotypes in the population (i.e. excluding the upper and lower 25th percentiles).

As expected, there was a strong linear correlation between the maximum and the interquartile mean (Fig 2). Our further expectation, supported by the above simulations, is that loci at which a mutator allele has existed in a population for a large number of generations can be identified from a characteristic pattern of variation, comprising a number of haplotypes with an unusually large number of derived alleles against a background of haplotypes with a typical number of derived alleles.

Diversifying selection is likely to increase both the interquartile mean and the maximum number of derived alleles. By contrast, a low-frequency mutator allele should affect only the maximum, and consequently we excluded from consideration windows with large values of the interquartile mean (>75th percentile). The remaining windows were arranged in order of a statistic *M*, defined as the residual distance to the regression line (the latter shown in red in Fig 2), with windows showing the greatest *M* ‐value considered as the best candidates for sites of ancient germline mutator alleles. We also removed candidate loci in which a high proportion of SNPs (> 5%) were not in Hardy-Weinberg equilibrium, as potentially indicative of artefacts. The top 20 genomic regions by *M* ‐value are shown in Table 1.

**Fig 2.**
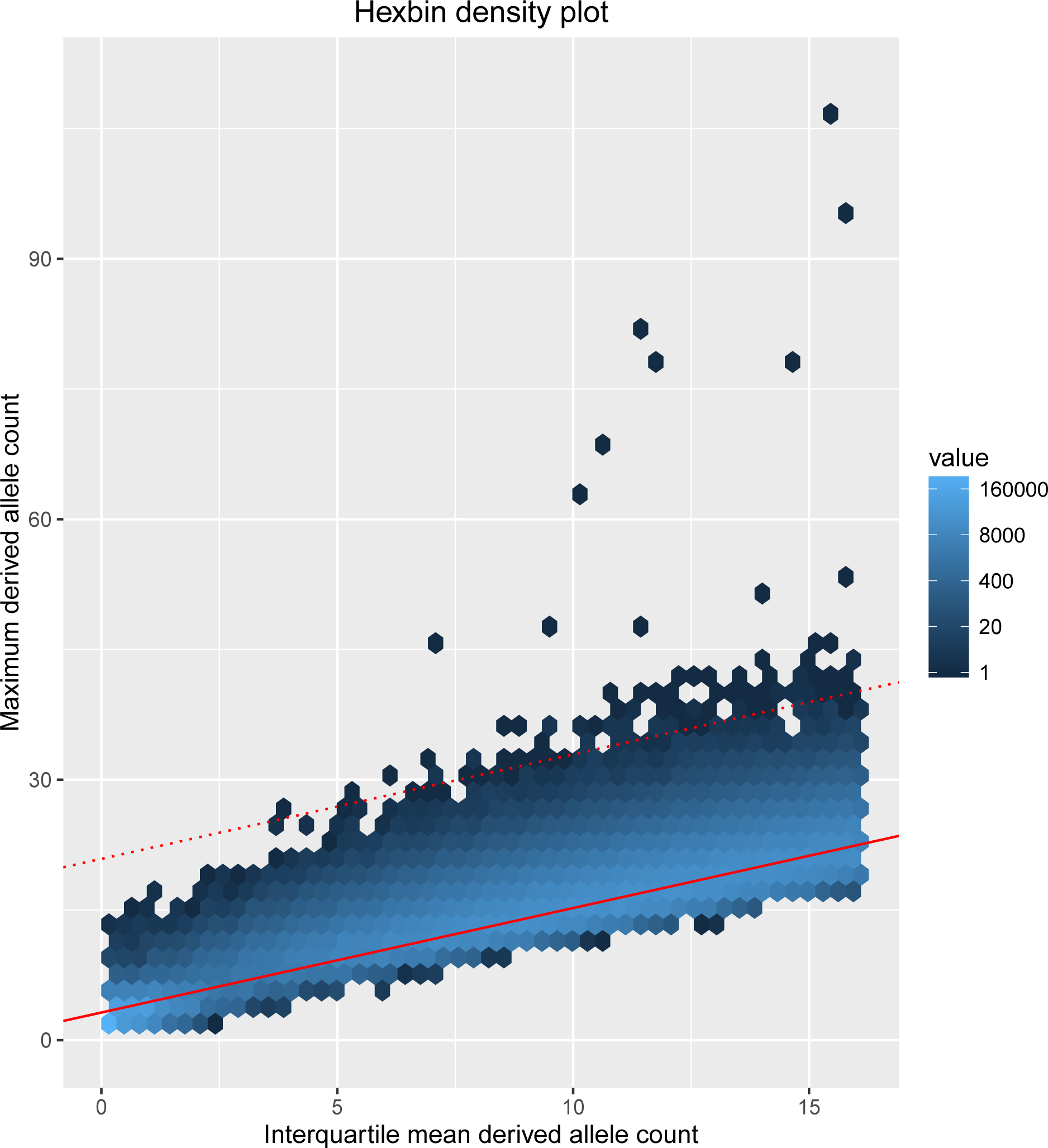
Relationship between the maximum and the interquartile mean number of derived alleles. A hexagon bin plot illustrating the relationship between the maximum and the interquartile mean number of derived alleles across all haplotypes in 10Kb windows from phase 3 of the G1K project. The data shown are for all populations combined, but the relationship is similar when populations are analyzed individually. The shading indicates the number of points within each hexagonal bin. The linear regression line is shown as a solid red line and the dashed red line identifies the candidate loci shown in Table 1. Note that each row of Table 1 can correspond to multiple points in the figure, due to multiple sliding windows overlapping the locus.

**Table 1.** Top 20 candidate mutator loci (relative to genome build hg19), ranked by residual.

| Chr | Position (Mb) | IQM | Max | Residual | Genes within 100 Kb |
| --- | --- | --- | --- | --- | --- |
| 1 | 202.4 | 15.4 | 106 | 84.28 | *UBE2T PPP1R12B* |
| 18 | 15.3 | 12.2 | 42 | 24.1 |  |
| 12 | 96.5 | 8.8 | 37 | 23.3 | *LTA4H ELK3* |
| 2 | 18.0 | 15.4 | 45 | 23.3 | *MSGN1 GEN1 KCNS3 SMC6* |
| 6 | 114.2 | 11.2 | 39 | 22.4 | *HDAC2 MARCKS* |
| 11 | 1.4 | 12.1 | 40 | 22.2 | *BRSK2 TOLLIP MOB2 TOLLIP-AS1* |
| 8 | 140.1 | 14.8 | 43 | 22.0 |  |
| 16 | 32.4 | 13.5 | 41 | 21.5 |  |
| 18 | 24.5 | 14.2 | 41 | 20.7 | *CHST9 AQP4* |
| 12 | 87.4 | 12.2 | 38 | 20.1 |  |
| 1 | 192.6 | 15.7 | 42 | 19.9 | *RGS1 MIR4426 RGS13* |
| 12 | 99.9 | 13.5 | 39 | 19.6 |  |
| 17 | 15.9 | 7.8 | 32 | 19.5 | *ADORA2B TTC19 NCOR1 ZSWIM7* |
| 14 | 90.4 | 15.3 | 41 | 19.4 | *EFCAB11 TDP1 KCNK13* |
| 6 | 130.9 | 11.9 | 36 | 18.6 |  |
| 8 | 42.0 | 11.2 | 35 | 18.3 | *KAT6A PLAT AP3M* |
| 10 | 25.0 | 14.6 | 39 | 18.2 | *ARHGAP21* |
| 21 | 24.1 | 7.2 | 30 | 18.1 |  |
| 12 | 44.6 | 7.2 | 30 | 18.1 |  |
| 5 | 17.3 | 4.0 | 26 | 18.1 | *BASP1* |

IQM: interquartile mean

To reduce the likelihood that divergent haplotypes that give rise to the entries in the table are the result mapping difficulties resulting from structural variants, we used the indel calls provided by the G1K project [30]. Two of the candidate loci in Table 1 (on chromosome 14 and at 192.6 Mb on chromosome 1) overlapped with a large structural variant (> 50bp). The structural variant on chromosome 14 is very rare, occurring in just 15 haplotypes from the G1K Phase 3 data, while the highly derived haplotype at this locus is much more frequent (occurring in 302 haplotypes). The opposite is the case for the structural variant at 192.6 Mb on chromosome 1, which is much more frequent (1871 haplotypes) than the highly derived haplotype at this locus (88 haplotypes). Therefore, in neither case is it plausible that mapping errors created by these structural variants are the cause of the observed signal. However, we cannot rule out that unreported structural variants are the cause of some of the candidate mutator loci that we identified.

### DNA repair genes are enriched at the candidate loci

Genes in proximity to the candidate loci shown in Table 1 are significantly enriched for just one cluster of overlapping biological processes, and encouragingly, these processes relate to DNA damage and repair. Five of these genes are annotated with the UniProt keywords *DNA damage* and *DNA repair* (p = 0.0004 and p = 0.0009, respectively), namely *GEN1*, *SMC6*, *TDP1*, *UBE2T* and *ZSWIM7*. Among these are genes that are annotated with the GO terms *double-strand break repair via homologous recombination* (GO:0000724; p = 0.005) and *DNA repair* (GO:0006281; p = 0.047). However, because genes are known to cluster on the genome by function, the above test of enrichment may be biased (for example, one candidate locus is responsible for two of the DNA-repair associated genes). To perform an unbiased test of enrichment of the candidate loci with DNA repair genes, we selected 20 loci at random and counted the number of DNA repair related genes within 100 Kb of each locus. Repeating this process 100,000 times, this number matched or exceeded the observed number 625 times (p = 0.006). This suggests that the enrichment of DNA repair genes is unlikely to be explained by functional clustering, and supports the existence of a subset of true positives among the list of candidate mutator alleles identified.

In addition to the known DNA repair genes, several other genes are plausible candidates for affecting the rate of germline mutation. *HDAC2* is a Class I histone deacetylase that has been shown to be enriched at replication forks [31] and to have a role in the DNA damage response, at least in the context of double strand break repair [32]. Mutations in *UBE2T* have recently been reported to cause Fanconi anemia [33], a condition characterised by genome instability and cancer susceptibility that frequently results from defective DNA repair genes. *BRSK2* is included among 296 genes in the PathCards DNA repair superpath [34] and recently *MOB2* has also been strongly implicated in the DNA damage response [35].

### Demographic simulations

We used demographic simulations to investigate the distribution of *M* ‐values in the absence of mutation rate polymorphism under different demographic scenarios (see Materials and Methods for details). As with the real data, the simulated data showed a strong linear relationship between the maximum (over haplotypes) derived allele count and the interquartile mean (Fig. S2). Two very different demographic scenarios were considered. In the first, a sample of size 2,500 (corresponding to the number of individuals in phase three of G1K) was taken from a single panmictic population. In the second, to explore the effects of population structure, the same number of individuals were sampled from a combined population resulting from pooling five separate populations that diverged 100 kyr ago. We found that both the relationship between interquartile mean and maximum number of derived alleles and the distribution of *M* ‐values were relatively insensitive to demography (Fig. S2). The simulations did not include selection and therefore generally had higher derived allele counts than the real data. Nonetheless, the most extreme value in the real data far exceeded the values found in the simulations (the most extreme *M* ‐values in the single population and multiple population simulation were 23.2 and 25.1, respectively). Although values as extreme as the remaining candidates in Table 1 were found in the simulated data, far more sliding windows exceeding the threshold for inclusion in Table 1 (residual ≥ 18) were found in the real data than in the simulated data (there were 95 such 10 Kb sliding windows in the real data, compared to 11 and 16 in the single and five population simulations, respectively). We also investigated the effect of mutation clustering caused by multinucleotide mutation events by adding these to the coalescent simulation (see Materials and Methods). This had limited impact on the results (the number of windows exceeding the threshold rose from 11 to 13 in the single population simulations).

Thus we conclude that the signals observed at loci listed in Table 1 are unlikely to have arisen through the demographic and neutral evolutionary processes represented in our simulations, and more likely to result from alternative processes such as inter-individual variation in mutation rate.

### Alternative explanations for the observed peaks

It is also necessary to consider other mechanisms, not included in our demographic simulations, that might have lead to excess divergence in a subset of haplotypes.

One example is a polymorphic gene conversion event affecting a subset of haplotypes, which could result in a block of divergent sequence. This would require gene conversion involving a segmentally duplicated region, in which the duplication of the target locus occurred prior to the divergence of humans from the other primates used to infer the ancestral state of genomic variants in this data. Under this scenario we may expect to find a subset of haplotypes that appear to be highly derived. A single gene conversion event giving rise to the diverged haplotypes would result in a phylogenetic tree of haplotypes consisting of one long internal branch separating two clades consisting of only short branches. The peak on chromosome 18 in Table 1 fits this description, but the remaining peaks do not (Fig. S3). For all of the remaining peaks, there is substantial diversity among the highly derived haplotypes, consistent with a greater rate of divergence than for the less highly derived haplotypes. Multiple gene conversion events would result in polyphyly of the diverged haplotypes, and this is not observed for most of the peaks; however, accurate phylogenetic inference may be exceptionally difficult in the gene conversion scenario.

Another possible mechanism might be an extreme multi-nucleotide mutation event, more substantial than those included in our simulations, giving rise to multiple mutations simultaneously. However, similarly to gene conversion, this would also create two clades, both consisting of short branches, separated by a long branch that reflects the multi nucleotide mutation. Thus for trees of this nature we cannot rule out either gene conversion or extreme multi-nucleotide mutation. However we note that such trees are also consistent with the effects of a mutator allele, since a relatively recent coalescence of the highly derived haplotypes would also produce a long internal branch separating clades consisting mostly of short branches. Indeed, this was frequently observed in the proof-of-concept simulations (Fig. S4).

Introgression of haplotypes derived from hybridisation between modern humans and Neanderthals as well as other ancient hominins has been reported [37–40]. However, this introgression is not likely to result in a subset of haplotypes with an excess of derived alleles. Alleles were designated as derived or ancestral by comparison to other primates and the number of derived alleles on a given Neanderthal haplotype should be similar to the number for a haplotype from modern humans. Moreover, almost all of the peaks in Table 1 are found in Africans, but only non-Africans have Neanderthal ancestry. The one exception is again the peak on chromosome 18. Indeed, for this peak we found that the highly derived haplotypes are enriched for haplotypes of likely Neanderthal origin (as determined by [41]). This peak is not close to any DNA repair genes and its removal from the list of candidates would in fact slightly increase the statistical significance of the association with DNA repair genes discussed above.

As a further potential alternative explanation, we also considered the case of a genomic segmental duplication segregating within the population, which could give rise to mapping errors in individuals carrying it, such that derived alleles called in the candidate locus are due to incorrectly mapped sequence reads. However, a simple duplication could cause at most an apparent doubling of divergence on affected haplotypes in this way, and therefore would be unlikely to be included as a candidate under the thresholds we have applied. A more complicated tandem duplication with several copies might give rise to higher apparent divergence, but such sites are very unlikely to have passed the filtering criteria used in the G1K calling protocol.

The peaks we observe in the derived allele count could potentially be caused by extremely strong selective pressure acting on a subset of populations from the G1K dataset. Such strong and localised selective pressure could cause a population or group of populations to have a large number of derived alleles within a genomic region, while the remainder of the populations in the dataset that did not experience the strong selective pressure have a typical number of derived alleles. Even if selective pressures strong enough to bring multiple linked derived alleles to high frequencies had acted on human populations we find no evidence for this hypothesis in the data, as, for most of the peaks, the highly derived haplotypes are found in human populations from multiple populations (Fig. S5).

Our proof-of-concept simulations suggest that most of the peaks in Table 1 are consistent with the effects of mutator alleles with large effect size which were maintained in the population for a large number of generations. The difference between the maximum and inter-quartile mean number of derived alleles in Table 1 is below 30 for all but the first locus, and in the low twenties for loci towards the bottom of the table. The proof-of-concept simulations also had large differences in the maximum and interquartile mean when the signal was detectable (just over 20 for simulations 8 and 9 in Fig. S1). The top locus in the table, however, has a much larger difference between the maximum and interquartile mean number of derived alleles. Although we could not find an alternative explanation for this signal, it seems difficult to reconcile with the effects of a mutator allele whose effect size would have enabled it to persist in the population for a large number of generations.

### Support from human trio data

Although we have considered the approach described here primarily as a means of detecting the remnant signatures of ancient mutator alleles, if such an allele were still active and linked to a highly-derived haplotype, then it might be possible to find supporting evidence in whole genome sequence data from trios. In principle, an active a mutator allele would give rise to an association between the presence of the highly-derived haplotype in the parents and an elevated number of *de novo* mutations in the offspring.

We investigated this possibility for the candidate loci shown in Table 1 by obtaining counts of *de novo* mutations in the offspring of complete trios and genome-wide parental genotype data from the G1K (n = 59) and GoNL (n = 248) projects [19, 20]. For each of the loci shown in Table 1 we tested for a positive association between the number of *de novo* mutations in the offspring and the presence in the corresponding parent of the highly derived haplotypes at the putative mutator locus (Table 2). P-values from two tests are reported for each peak, one based on fitting a robust linear model and the second based on permutation (referred to as *p_t_* and *p_perm_* in Table 2, respectively; see Materials and Methods for details). The analysis was carried out separately for male and female parents and for the two projects, to avoid biases resulting from differences in study methodology confounded with differences in populations of origin. In each case we required at least five parents with a highly derived haplotype to perform the test.

**Table 2.** P-values (*p_t_, p_perm_*) from tests of elevated rate of *de novo* mutation in trios.

| Chr | Position (Mb) | G1K Paternal | G1K Maternal | GoNL Paternal | GoNL Maternal |
| --- | --- | --- | --- | --- | --- |
| 1 | 202.4 | NA | NA | NA | NA |
| 18 | 15.3 | NA | NA | NA | NA |
| 12 | 96.5 | NA | NA | NA | NA |
| 2 | 18 | 0.49, 0.48 | 0.86, 0.88 | NA | 0.26, 0.26 |
| 6 | 114.2 | NA | NA | NA | NA |
| 11 | 1.4 | 0.20, 0.21 | 0.0033, 0.0039 | 0.95, 0.95 | 0.91, 0.91 |
| 8 | 140.1 | 0.75, 0.75 | 0.29, 0.30 | 0.16, 0.16 | 0.13, 0.13 |
| 16 | 32.4 | NA | 0.034, 0.067 | NA | NA |
| 18 | 24.5 | NA | NA | NA | NA |
| 12 | 87.4 | NA | NA | 0.49, 0.48 | 0.41, 0.41 |
| 1 | 192.6 | 0.049, 0.064 | 0.24, 0.26 | NA | NA |
| 12 | 99.9 | NA | NA | NA | NA |
| 17 | 15.9 | NA | NA | NA | NA |
| 14 | 90.4 | 0.63, 0.62 | 0.55, 0.55 | 0.75, 0.76 | 0.96, 0.97 |
| 6 | 130.9 | NA | NA | NA | NA |
| 8 | 42.0 | 0.26, 0.26 | 0.59, 0.59 | NA | NA |
| 10 | 25.0 | 0.49, 0.21 | 0.56, 0.57 | 0.45, 0.44 | 0.13, 0.13 |
| 21 | 24.1 | 0.67, 0.68 | 0.51, 0.51 | NA | NA |
| 12 | 44.6 | 0.024, 0.040 | 0.44, 0.44 | 0.43, 0.42 | 0.10, 0.10 |
| 5 | 17.3 | 0.22, 0.23 | 0.70, 0.70 | 1.0, 1.0 | 0.87, 0.87 |

In the case of the G1K dataset 21 tests were performed (10 paternal and 11 maternal). Of these, two showed a nominally significant positive association between the number of *de novo* mutations and the presence of the highly derived haplotype in the parent in both statistical tests (Table 2). The most significant of these was for the peak just upstream of *BRSK2* on chromosome 11 (illustrated in Fig 3). With the GoNL dataset, 15 tests could be performed (seven paternal and eight maternal), but no significant associations were observed (Table 2). Considering that 36 tests in total were carried out in total, the associations in Table 2 do not remain significant following Bonferroni correction, and additional diverse trio datasets may be necessary to confirm these associations definitively. Data forming the basis for the tests shown in Table 2 are provided as S1 Table and S2 Table.

Paternal age at conception is strongly correlated with the number of *de novo* mutations in offspring [8]; however, paternal age was not available for the G1K data, while for the GoNL data we included the ages of both parents at conception in the linear model relating *de novo* mutation count to the presence of the highly derived haplotype. There remained no significant positive associations in the case of this dataset. For the G1K dataset population of origin information was available for the samples; however, there was no statistically significant relationship between population of origin and the *de novo* mutation count (Fig. S6).

Interestingly, the candidate locus that shows the strongest evidence of an association with the *de novo* mutation count in the G1K trios is just upstream of *BRSK2* (Fig 3b), a highly conserved serine threonine kinase which is preferentially expressed in brain and testis and shows enhanced activity in response to DNA damage [42, 43]. The observed association was *maternal* rather than paternal; however, expression in testis is consistent with a role in germline DNA replication.

At the same time, this haplotype is not associated with an elevated *de novo* mutation count in the GoNL data (Table 2). A possible explanation for this is that despite its much larger sample size, the GoNL cohort, which comprises individuals of Dutch ancestry, is substantially less diverse than the G1K cohort. As noted above, although the highly derived haplotype was observed in parents of trios in the GoNL cohort, its presence does not necessarily imply the presence of the causative mutator allele: a recombination event on the ancestral lineage leading to the highly derived haplotype in the Dutch population may have separated the two. Indeed, in this way it is possible for an extinct mutator allele to leave a fossil peak in the derived allele count of a contemporary population, and it is plausible that many of the signals we have detected are of this nature.

The candidate mutator locus on chromosome 14 is also noteworthy. This peak overlaps precisely with an exon of the tyrosyl-DNA phosphodiesterase 1 (*TDP1*) gene (Fig. S7). The product of *TDP1* has a role in the repair of stalled topoisomerase I-DNA complexes. This candidate is not supported by the available trio datasets, despite the highly derived haplotypes being found in sufficient numbers in both cohorts, suggesting that its occurrence on the *TDP1* gene is either a striking coincidence or that it may be a fossilised peak resulting from a mutator allele that is extinct or at least much rarer than the highly derived haplotypes to which it has given rise.

**Fig 3.**
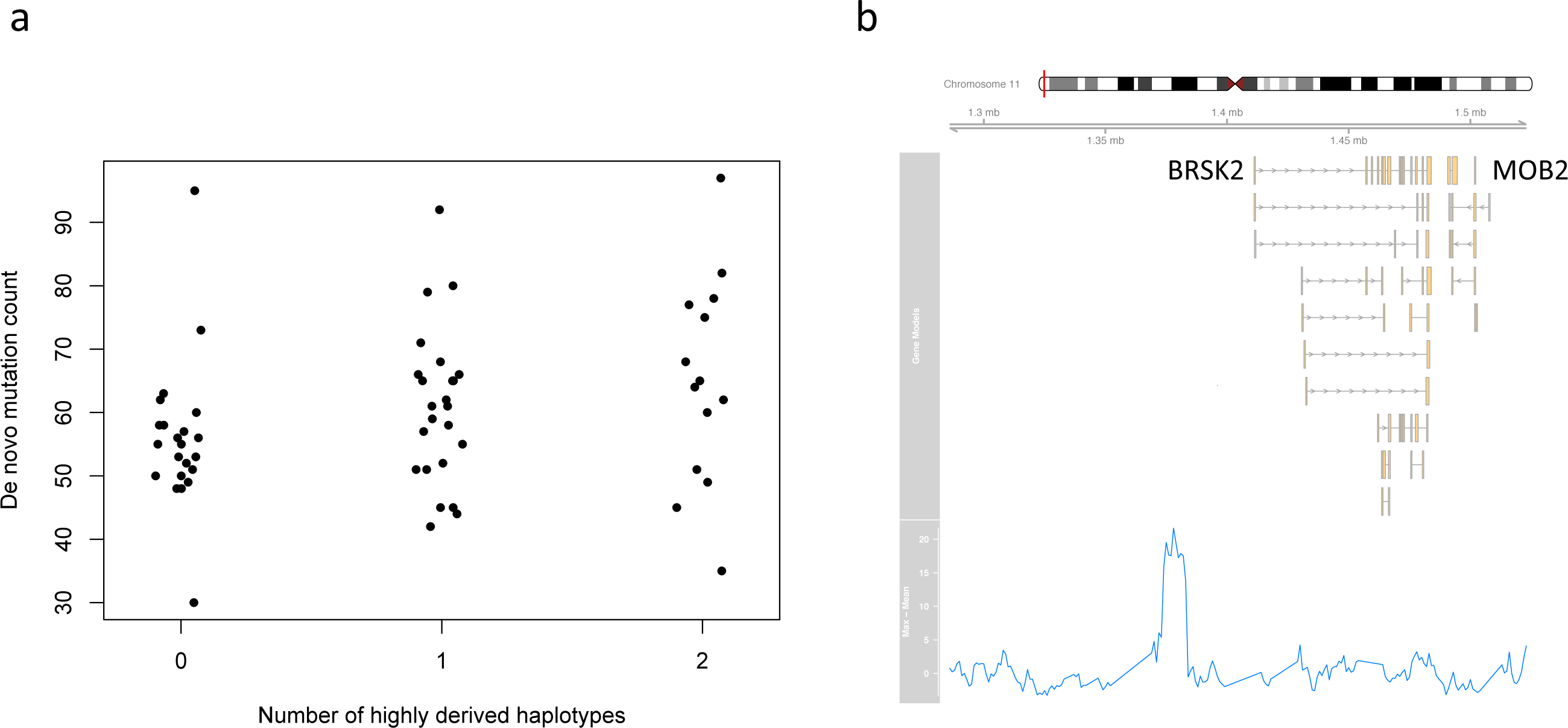
Putative mutator allele on chromosome 11. Relationship between the number of *de novo* mutations in the offspring and the maternal number of highly derived haplotypes of the putative mutator allele on chromosome 11 (a). Location of the putative mutator locus on chromosome 11, defined as a peak in the difference between the maximum and interquartile mean number of derived alleles across haplotypes (b).

## Conclusions

We have shown that when a genomic locus contains a polymorphism in which one allele increases the germline mutation rate, a characteristic signature may result, comprising a subset of haplotypes that are locally more divergent than other haplotypes at that locus. Whether this signature can be detected depends on the magnitude of the effect and on the time since the mutator allele arose. The number of additional mutations on an unrecombined segment of expected length carrying the mutator allele is independent of the time since the mutator allele arose, but the likelihood of detecting a fixed number of additional mutations over the background increases as the segment length decreases and is thus higher for older mutations.

We searched for such signatures in data from Phase 3 of the 1000 Genomes Project, and found candidate human genetic loci that may have once contained (and in some cases may still contain) germline mutator alleles. Consistent with this expectation, the set of genes located in the vicinity of these loci is enriched for genes involved in DNA repair. A test for the continued presence of active mutator alleles in these haplotypes, by looking for association with high numbers of *de novo* mutations passed to offspring in two trio sequencing cohorts, found support for association in two cases, but no associations with overall significance.

It may therefore be that mutator alleles are no longer present, or are now very rare, in most of the candidate loci we identified. However the trio cohorts we examined were small relative to what may be feasible in future studies, and in one case (the GoNL data) represented a population which is comparatively homogeneous in terms of its genetic ancestry. Further trio cohorts from more genetically diverse populations may be necessary to test some of the candidates we identified. Such studies will also shed light on the full extent of polymorphism in the rate of *de novo* mutation in human populations.

It is difficult to conclusively exclude alternative explanations for the signals we have detected. However we have considered several such explanations and have shown that they are either far less likely to generate signals of this nature (in the case of neutral demographic processes) or would generate loci with characteristics not seen in these cases (for example in the case of a mis-mapped segmental duplication).

More generally, it is far from implausible *a priori* that there are genetic factors affecting the germline mutation rate in humans, and that there may arise from time to time mutator alleles with a moderately large effect size. We have also shown that such alleles can plausibly persist for many generations and leave a detectable signal that survives after they have disappeared. It may also be that different such alleles affect the mutation process in different ways, and that as well as changing the overall germline mutation rate they give rise to different mutational signatures. Indeed recent studies have found variation in the mutational spectrum (expressed in terms of the relative frequency of different triplet sequence contexts for de novo mutations) between human populations and over time, including evidence for past activity of mutator alleles that are now rare or extinct ( [18, 54]).

On longer timescales, comparison of mutation rates estimated from inter-specific divergence (e.g. [44]) with measurements of the mutation rate in contemporary populations points to a slowdown in human and great ape evolution [55]. Some of this may be due to changes in life history parameters such as generation time; however, there may also have been changes in the underlying per-generation mutation rate, or more specifically, changes in cellular mutation rates during development and gametogenesis, perhaps driven by genetic variation at loci such as those detected here. Indeed, it is worth considering that the higher mutation rate inferred for some haplotypes at candidate loci may represent the ancestral rather than derived state, such that the mutations in question resulted in anti-mutator rather than mutator alleles.

## Materials and Methods

### Proof of concept simulations

We used a coalescent approach to simulate the effects of a mutation that increases the germline mutation rate. The mutator allele is introduced at an initial frequency *p*_0_ at time *t*_0_ and we assume no mutation between the mutator allele and wild-type after time *t*_0_. We use a constant effective population size of 10,000 individuals and incorporate recombination, occurring at a constant rate across the simulated genomic region (see below for simulations of null models with variable recombination rate and demographics). The mutator allele is associated with a selective coefficient, *s*, and we incorporate selection against the mutator allele using an approach described by Kaplan *et al.* [46]. We first obtained the deterministic allele frequency trajectory in the forward direction, with *s* = 0.002. Because we require that chromosomes carrying the mutator allele have an elevated mutation rate the simulations could not be performed using existing implementations of the coalescent with recombination and selection (e.g. [47]) and were instead implemented as a Perl script, which is available from the authors, on request. To allow for the effects of the mutator allele on the rate of mutation of the homologous chromosome in heterozygous individuals, at each generation the mutation rate was elevated for chromosomes that did not carry the mutator allele with a probability equal to the frequency of the mutator allele in that generation.

### Demographic simulations

To examine the genome-wide distribution of maximum and inter-quartile mean number of derived alleles per window, we simulated populations evolving under a model of neutral evolution and without mutation rate polymorphism. Two simulations of 5000 haploid whole genome sequences were carried out, one comprising a single population with constant effective population size, and another comprising five separate populations diverging 100 kyr ago, to explore the potential effects of population structure on the signal of interest. All populations simulated were of equal size with a scaled mutation rate *θ* = 0.001, matching the mean genome-wide heterozygosity in modern human populations. Simulations were carried out using the coalescent simulator msprime [48], with a variable recombination rate matching that found in the human genome (including recombination hotspots) [49], and a uniform genome-wide mutation rate.

### Genome scan

Simulations suggested that a mutation that increases the rate of germline mutation and is maintained in a population for a sufficiently long time can result in a genomic region in which haplotypes linked to the mutator allele have a large number of derived alleles and the remaining haplotypes have typical numbers of derived alleles (i.e. the effect of the mutator allele on unlinked haplotypes is the same as its effect on the rest of the genome). Variant calls in .vcf format were downloaded for phase 3 (release v5a) of the 1,000 Genomes project [20]. Putative ancestral alleles (derived by the 1,000 Genomes project from Ensembl Compara release 59 [29]) were obtained from the vcf files, restricting to SNPs with high confidence ancestral alleles. Using a sliding window (window size: 10 Kb; step size: 1 Kb) we calculated the interquartile mean (also called the 25% trimmed mean) and the maximum number of derived alleles in each window. There was a linear relationship between the maximum and the interquartile mean, reflecting the fact that windows within which the interquartile mean number of derived alleles was large also had a large value for the maximum number of derived alleles (over all haplotypes). We fitted linear regression models to the maximum, as a function of the interquartile mean and calculated the distance from the regression line for each window. Windows with an elevated value of the interquartile mean (>75th percentile) were excluded and the remaining windows were sorted by distance to the regression line. *χ*^2^ tests of Hardy-Weinberg equilibrium (at *alpha* = 0.05) were carried out separately within each population for each SNP within the candidate regions. Windows within which a high proportion (>5%) of SNPs rejected Hardy-Weinberg equilibrium were removed. This resulted in the removal of nine loci.

### Trio validation

We obtained *de novo* mutation counts in offspring and parent variant call data for human parent-offspring trios from the G1K (n = 59) and GoNL (n = 248) projects [19, 20]. For each candidate mutator locus we determined the number (0, 1 or 2) of copies of the highly derived haplotype in each parent. For the phased G1k data we plotted the number of derived alleles per haplotype and if this distribution included a subset of highly derived haplotypes we selected a threshold that gave the best separation between the highly derived and typically derived haplotypes (Fig. S8) and used this to classify each haplotype. For unphased data (GoNL) we plotted the number of derived alleles per individual. When the highly derived haplotypes occurred at low frequency this distribution was typically bimodal (all individuals had 0 or 1 highly derived haplotype), becoming trimodal when the highly derived haplotypes occurred at higher frequency (corresponding to individuals with 0, 1 or 2 highly derived haplotypes). In each case we set a threshold number of derived alleles that gave the best separation of the peaks in the distribution of the derived allele count (Fig. S9). We then tested for an association between the offspring *de novo* mutation count and the number of copies of the highly derived haplotype by fitting robust linear regression models using the rlm function from the MASS package [50] in R [51], with the default M-estimation method. The G1K and GoNL trios were analysed separately to avoid the potential for confounding between study methodologies and population of origin and results are reported separately for each parent (to enable the detection of parent-of-origin specific effects). Parental age at conception was available for the GoNL trios and included as a covariate in the linear models. P-values for the null hypothesis that the regression coefficient for the offspring *de novo* mutation count as a function of the number of copies of the highly derived haplotype was not greater than zero were calculated in two ways: from the right tail of the distribution of the regression coefficient t-statistic (referred to as *p_t_* in Table 2) and by permutation. For the latter we performed 10,000 permutations of the *de novo* mutation counts and determined the proportion of times the t-statistic from the robust regression for the permuted data was greater than or equal to the value for the unpermuted data (reported as *p_perm_* in Table 2). Correction for multiple testing was carried out using the Bonferroni method.

### Enrichment of DNA repair genes close to candidate loci

We identified all genes within 100 Kb of each of the candidate mutator loci (excluding pseudogenes) and tested for functional enrichment using DAVID [52, 53], version 6.8 with default parameters (on 3/11/2016). Given that there were several genes found within 100 Kb of many of the candidate mutator loci the phenomenon of functional gene clustering means that this set of genes is not independent, creating a potential for bias in the statistical tests of enrichment. We therefore also devised a test based on randomisation of the genomic loci. For this test 20 loci (equal to the number of loci in Table 1) were sampled at random from the regions of the data for which we had variant call data and the number of DNA repair genes (annotated with GO term GO:0006281 in Ensembl 84) within 100 Kb of these random loci was determined. This was repeated 100,000 times and the numbers of DNA repair genes in the random data was compared to the observed number of genes with this GO term within 100 Kb of the candidate mutator loci.

## Acknowledgments

This study makes use of data generated by the Genome of the Netherlands Project and the 1000 Genomes Project. We thank the scientists and participants of the 1000 Genomes and Genome of the Netherlands projects for generously sharing their data.

## Supporting Information

**Fig. S1 Proof of concept simulations**. Ten random simulations, equivalent to the simulation shown in Fig. 1.

**Fig. S2 Demographic simulations**. (a,b) Hexagon bin plots illustrating the relationship between the maximum and the interquartile mean number of derived alleles across all haplotypes in 10Kb windows for two simulated demographic scenarios, consisting of a single population (a) or five populations separated 100 kya (b). The shading indicates the number of points within each hexagonal bin. The linear regression line is shown as a solid red line and the dashed red line corresponds to the threshold for the candidate loci shown in Table 1. (c) Quantile-quantile plot of the residuals from the regression line in the single population and five-population simulations.

**Fig. S3 Plots of the 20 candidate mutator loci shown in Table 1**. For each candidate mutator locus in Table 1 we show a phylogenetic tree (left), constructed from a 10 Kb window overlapping the peak. Due to the difficulty displaying all 5008 haplotypes from the 1000 Genomes Project we sampled 100 haplotypes at random and supplemented these with up to approximately 50 of the highly derived haplotypes, again at random. Bottom right panels illustrate for this reduced set of haplotypes the number of derived (red) and ancestral alleles (blue) in each haplotype across the 10Kb window. Each haplotype is represented as a row on the plot and the haplotypes with the largest number of derived alleles are grouped at the top of the plot. Tick marks indicate the positions of individual SNPs. The top right panels show the maximum (red) and interquartile mean (green) number of derived alleles, as well as the number of derived alleles on actual haplotypes (grey).

**Fig. S4 Trees from proof-of-concept simulations**. Trees from the subset of the ten proof-of-concept simulations that showed a peak in the maximal derived allele count.

**Fig. S5 Population of origin of the highly derived haplotypes**. The proportion of the highly derived haplotypes at each peak corresponding to populations from Africa, Asia, America and Europe. The rightmost bar shows the proportions of each continent of origin among all G1K phase 3 samples.

**Fig. S6 Population of origin effect on *de novo* mutation count**. Stripchart showing *de novo* mutation count as a function of population of origin in the G1K data.

**Fig. S7** Candidate mutator locus within the *TDP1* gene.

**Fig. S8** Derived allele count thresholds used to decide highly derived haplotypes for the G1K data.

**Fig. S9** Derived allele count thresholds used to decide the number of highly derived haplotypes in each individual for the unphased GoNL data

**S1 Table. Trio data from 1000 Genomes**. The number of de novo mutations and number of copies of the highly derived haplotype in trios from the 1000 Genomes Project.

**S2 Table. Trio data from the Genome of the Netherlands Project**. The number of de novo mutations and estimated number of copies of the highly derived haplotype in trios from the Genome of the Netherlands Project.

